# Optimization of breeding conditions in *Danionella cerebrum* to advance its use as a vertebrate model

**DOI:** 10.64898/2026.10.10.758160

**Authors:** Chih Chi Kao, Koji Ando, Jason A. da Silva, Teruhisa Kawamura, Kazu Kikuchi

## Abstract

We systematically evaluated tank configurations and spawning-tube parameters to improve egg production and collection efficiency in *Danionella cerebrum*. A shallow breeding tank with a black bottom significantly increased egg production while facilitating egg collection. Further optimization of spawning-tube parameters revealed a strong effect of tube diameter on reproductive performance, with narrower tubes improving fertilization efficiency and a 4-mm inner diameter yielding the highest egg production. Tube length also influenced breeding performance and handling efficiency. Longer tubes tended to increase egg production, whereas 5-cm tubes were more convenient for routine egg collection. The greater water volume of 10-cm tubes also enabled short-term in-tube embryo culture, potentially improving the efficiency of germline screening for transgenic founders. Notably, increasing the number of available spawning tubes did not further enhance egg production, leading us to examine patterns of spawning-tube usage. This analysis revealed a previously unrecognized preference of *D. cerebrum* for spawning sites previously used by other breeding pairs. Together, these findings establish practical breeding parameters that improve reproductive output and streamline egg collection and embryo handling in *D. cerebrum*.

**Summary Statement:** We improved breeding conditions for the tiny transparent fish *Danionella cerebrum*, identifying tank and tube designs that increase egg production and simplify egg collection, embryo handling, and laboratory breeding.

## Introduction

*Danionella* represents a unique group of miniature teleosts notable for their exceptional transparency and minimal pigmentation (Roberts, 1986). Of these species, *Danionella cerebrum* has been most extensively established as a laboratory model and has emerged as a promising vertebrate system for *in vivo* imaging in neurobehavioral research (Schulze et al., 2018). Unlike zebrafish and other model organisms, *D. cerebrum* remains fully transparent throughout its life, lacking scales and possessing entirely translucent skin and internal organs. In addition, *D. cerebrum* lacks a cranial bone at the dorsal surface of the brain (Schulze et al., 2018), likely due to a developmental truncation (Britz and Conway, 2016). This unique anatomical trait, together with its exceptionally small adult brain and rich behavioral repertoire (Schulze et al., 2018), establishes *D. cerebrum* as an outstanding model for brain imaging and behavioral research (Conway et al., 2021; Akbari et al., 2022; Lindemann et al., 2022; Rajan et al., 2022b; Tatarsky et al., 2022; Lee and Briggman, 2023; Bass and Perelmuter, 2024; Cook et al., 2024; Vasconcelos et al., 2024; Veith et al., 2024; Zada et al., 2024; Fouke et al., 2025; Penalva-Tena et al., 2025).

Importantly, *D. cerebrum* is genetically tractable and readily supports modern genetic technologies. The availability of a reference genome (Kadobianskyi et al., 2019) enables efficient CRISPR/Cas9-mediated gene disruption (Schulze et al., 2018) and provides a foundation for the future implementation of targeted knock-in strategies to precisely insert exogenous DNA into defined genomic loci. In addition, as in zebrafish and other teleost model organisms, stable transgenic lines can be generated by microinjecting plasmid constructs into one-cell-stage embryos (Schulze et al., 2018). Together, these approaches facilitate the generation of fluorescent reporters and biosensors for high-resolution visualization of cellular and molecular dynamics *in vivo*, as well as genetic tools for functional interrogation of the underlying biological processes. The combination of genetic tractability and exceptional lifelong optical transparency positions *D. cerebrum* as a powerful vertebrate model for real-time, whole-organism imaging and extends its utility beyond neurobiology to the investigation of diverse biological processes across a wide range of biomedical research fields.

Despite this considerable potential, several practical limitations currently constrain the widespread adoption of *D. cerebrum*. Compared with other established model organisms, *D. cerebrum* currently lacks the extensive resources available for more mature experimental systems, including comprehensive epigenomic and transcriptomic datasets, large collections of transgenic and mutant lines, validated molecular reagents such as antibodies, and centralized community resources. As the *D. cerebrum* research community continues to grow, these resources are expected to expand substantially, with several initiatives already underway, including the generation of a genome assembly and annotation (Kadobianskyi et al., 2019), as well as a single-cell transcriptomic atlas (Atabay et al., 2026). Consequently, many of the current technical limitations are likely to diminish over time, further broadening the utility of this model across a wide range of research applications (Lam, 2022; Atabay et al., 2026).

By contrast, the relatively low fecundity of *D. cerebrum* represents an intrinsic constraint that is unlikely to be manipulated substantially and therefore remains the principal bottleneck to its broader application. Zebrafish routinely produce several hundreds of eggs from a single spawning (Streisinger et al., 1981), and medaka spawn daily, producing 10–30 eggs under standard laboratory conditions (Kinoshita et al., 2009). However, *D. cerebrum* is characterized by infrequent and irregular spawning, with a small clutch size of approximately 12 eggs (Rajan et al., 2022a), limiting the scale and throughput of experiments, including the generation of genetically modified lines. In this study, we systematically evaluated environmental and spawning substrate parameters affecting egg production efficiency and fertilization success in *D. cerebrum*. Based on these findings, we propose practical breeding procedures that improve routine egg recovery and facilitate the screening of transgenic lines, thereby enhancing the utility of *D. cerebrum* as a vertebrate model for biomedical research.

## Materials and Methods

### Tank specifications

The aquarium tanks used in this study were custom manufactured by IWAKI (Tokyo, Japan). The standard housing tank had a capacity of 1.2 L, measured 80 × 150 × 230 mm (W × H × D), and was constructed entirely of transparent polycarbonate. The wide-and-shallow tank had a capacity of 3 L, measured 300 × 100 × 250 mm (W × H × D), and was constructed either entirely of transparent acrylic or with transparent acrylic walls and a matte black bottom panel.

### D. cerebrum husbandry

A breeding colony of *D. cerebrum* was established from adult fish provided by Dr. Benjamin Judkewitz, Charité–Universitätsmedizin Berlin, Germany. Fish were maintained as previously described (Rajan et al., 2022a), with minor modifications detailed below. Briefly, fish were housed in an IWAKI recirculating aquaculture system (IWAKI, Tokyo, Japan) under a 14:10 h light photoperiod at 27 °C, with water maintained at pH 7.3 and a conductivity of 350 µS cm⁻¹. LED lights positioned directly above the tanks were turned on for approximately 1 h during egg collection. All animal experiments were conducted in accordance with institutional and national guidelines for the care and use of laboratory animals and were approved by the Animal Ethics Committee of the National Cerebral and Cardiovascular Center.

### Live feed culture

*Paramecium* and *Artemia* (Kitamura, Kyoto, Japan) were cultured and prepared for feeding according to standard protocols (Westerfield, 2000). Freshwater rotifers (*Brachionus*) were cultured and prepared with modifications to a previously published protocol (Aoyama et al., 2015), as detailed below.

The procedure consisted of three phases: (i) rotifer egg hatching and continuous culture, (ii) the daily rotifer production cycle, and (iii) rotifer harvesting and feeding. For rotifer egg hatching and continuous culture, a capsule of dried rotifer eggs (Sakura Medaka, Kobe, Japan) was added to 3 L of aquarium water in an Erlenmeyer flask (010530-3000; SIBATA Scientific Technology, Saitama, Japan). The flask was loosely covered with aluminum foil and incubated at 28 °C in a temperature-controlled incubator (MIR-254-PJ; PHC, Tokyo, Japan) for 3–5 days until hatching. Once hatching was confirmed, half of the culture (1.5 L) was transferred to a new flask to establish a continuous culture, which was maintained by daily replenishment with 1.5 L of fresh aquarium water. The remaining half (1.5 L) was used to initiate the daily rotifer production cycle described below. Additional flasks were prepared as needed.

For the daily rotifer production cycle, the culture (1.5 L) was replenished with 1.5 L of fresh aquarium water containing 2 mL of *Chlorella* (Chlorella-V12; Chlorella Industry, Fukuoka, Japan), continuously aerated through an air stone connected to a NON-NOISE W1000 air pump (Japan Pet Design, Tokyo, Japan), and incubated overnight under the same conditions. The following day, half of the culture (1.5 L) was harvested, and the remaining culture was replenished with 1.5 L of fresh aquarium water containing 2 mL of *Chlorella* to maintain the culture.

For rotifer harvesting and feeding, the harvested culture (1.5 L) was passed through a 40 µm sieve. Rotifers retained on the sieve were rinsed into a 200 mL beaker with reverse osmosis water, adjusted to a final volume of 150 mL with aquarium water, and dispensed into embryo tanks using a disposable plastic pipette. This harvest-and-replenishment cycle was repeated daily.

### Rearing and feeding of *D. cerebrum*

Following mating, *D. cerebrum* embryos were collected and incubated in E3 medium (5 mM NaCl, 0.17 mM KCl, 0.33 mM CaCl_2_, and 0.33 mM MgSO_4_) supplemented with 10^−5^% methylene blue in 85 mm Petri dishes (AS ONE, Osaka, Japan) at room temperature (25 °C) for 5 days, during which dead embryos and debris were removed daily and the E3 medium was replaced as necessary until 3 days post-fertilization (dpf). On 5-dpf, the larvae were transferred to 1.2 L housing tanks containing approximately 1 L of aquarium water and fed live rotifers once daily until 8-dpf, followed by rotifers and *Artemia* until 14-dpf.

Between 5-dpf and 14-dpf, approximately 0.5 mL of fresh E3 medium containing *Paramecium* was added daily into the tank. Although *Paramecium* was not readily consumed by *D. cerebrum* larvae, it scavenged food debris and other organic material, allowing the larvae to be maintained without routine water changes or tank cleaning, requiring only the daily addition of fresh medium. From day 14 onward, the fish were transferred to a recirculating aquarium system and fed *Artemia* once daily in the morning for routine maintenance colonies or twice daily, in the morning and afternoon, for breeding colonies.

### Imaging and microscopes

Sex determination and fertilization assessment were performed using an SZX7 stereomicroscope (Olympus, Tokyo, Japan). Images of multiple egg clutches and adult females within the silicone tubes, as well as lateral views of the tubes, were acquired using an HC-V360M digital video camera (Panasonic, Osaka, Japan). Cross-sectional images of the silicone tubes and images of adult male and female *D. cerebrum* were acquired using an S9D stereomicroscope equipped with a Flexacam C5 camera (Leica Microsystems, Wetzlar, Germany).

### Sex identification

Sex was determined by visual inspection based on the position of the anus, as previously described (Britz et al., 2021). Briefly, fish were examined under a stereomicroscope at 3 months of age and identified as males if the anus was located between the pelvic fins, or as females if it was positioned posterior to the pelvic fins and closer to the anal fin. Females could also be readily distinguished by the presence of developing or mature oocytes visible through the transparent body wall.

### Reproduction assays

To assess reproductive performance in *D. cerebrum*, we used adult males and females aged 9–15 months. Comparable age and sex distributions were maintained across all experimental groups. Semi-transparent Laboran silicone tubes (AS ONE, Osaka, Japan) of the indicated diameters and lengths were used as spawning substrates (Schulze et al., 2018; Rajan et al., 2022a) and were provided at a tube-to-fish ratio of 1:2 unless otherwise indicated. Fish were acclimatized for the period indicated in each experimental condition, during which spawned eggs were collected daily and fertilization was assessed under a stereomicroscope approximately 24 hours after egg collection. Following the acclimatization period, fish were redistributed between the high- and low-performing groups to balance baseline reproductive performance across the experimental groups as closely as possible in selected experiments. The experimental period was then initiated under the indicated conditions, with egg production and fertilized egg number assessed daily as described above.

### Spawning tube usage analysis

Spawning tube preference during mating was assessed using 15-cm-long silicone tubes with a 4-mm inner diameter. 15-cm tubes were used in this assay to maximize clutch retention and thereby enable accurate quantification of egg numbers within each tube. Each tube was manually embossed with a unique pattern using a P1000 pipette tip (Watson, Kobe, Japan), producing subtle markings that were readily identifiable under a stereomicroscope but minimally visible to the naked eye. Three tubes bearing unique embossed patterns were then placed in each flat-B tank containing 20 fish, with an equal female-to-male ratio. The tubes were retrieved from the tanks and sealed at one end with commercially available rubber stoppers (Uxcell, Shenzhen, China), with the water retained within the tubes by surface tension. The numbers of clutches and eggs were counted daily for 10 days. Days on which no or only a single clutch was produced were excluded because repeated use of the same tube could not be assessed.

### In-tube embryo rearing

Multiple 10-cm-long silicone tubes with a 4-mm inner diameter were placed in a flat tank containing 20 fish at a 1:1 female-to-male ratio, and the fish were allowed to spawn for 24 h. The tubes were retrieved from the flat-B tanks, sealed at one end with rubber stoppers, and incubated at room temperature for 3 days. Following in-tube rearing for 3 days, embryos were transferred to 1.2-L housing tanks containing approximately 1.0 L of aquarium water at a density of 9–13 fish per housing tank for subsequent developmental assessment. Embryos were anesthetized with 0.01% tricaine (Sigma-Aldrich, St. Louis, MO, USA), embedded in 2.5% methylcellulose (Sigma-Aldrich, St. Louis, MO, USA), and imaged using a THUNDER Imager (Leica Microsystems, Wetzlar, Germany) to evaluate for developmental abnormalities.

## Results

### Optimization of tank configurations

*D. cerebrum* is known to spawn eggs in clutches within narrow crevices (Schulze et al., 2018). Under laboratory conditions, silicone tubes mimic these natural spawning sites, allowing breeding to occur directly in housing tanks without transferring fish to dedicated breeding tanks (Schulze et al., 2018; Rajan et al., 2022a). To examine whether tank design influences reproductive performance, we developed a flat tank (Fig. 1A), a custom housing tank designed to facilitate both breeding and routine egg collection. In this design, the water depth was reduced from 10-cm in the standard housing tank (hereafter referred to as the housing tank) to approximately 4-cm (Fig. 1A), based on previous reports that zebrafish and medaka preferentially spawn in shallow water (Sessa et al., 2008; Kamide et al., 2020). The shallower water was also expected to facilitate the recovery of spawned eggs using a standard disposable plastic transfer pipette (Fig. 1A). In addition, the bottom area was increased from 94.5 cm² in the housing tank to 696 cm² to provide greater horizontal space for courtship and spawning (Fig. 1A).

**Figure 1.**
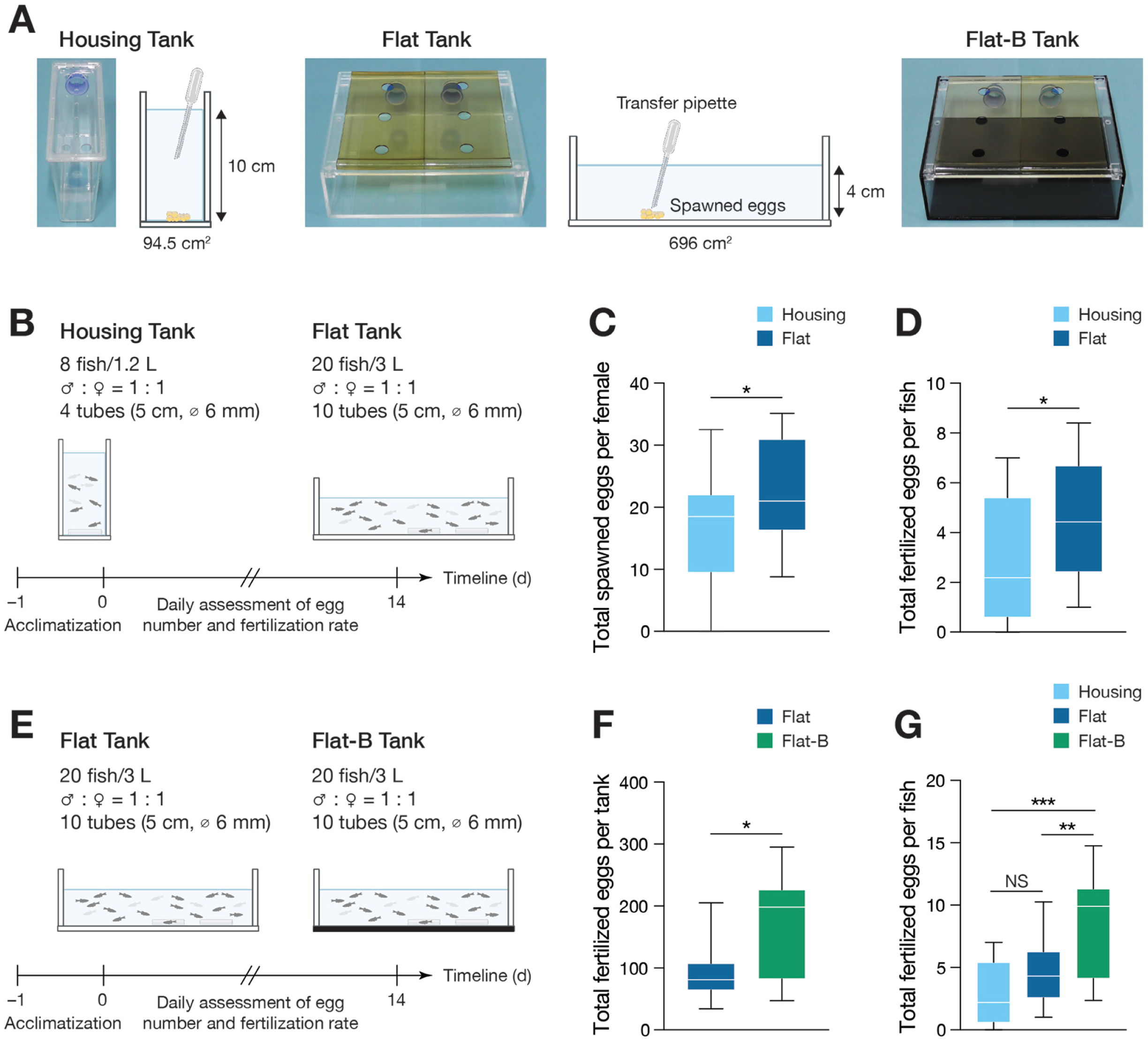
Reproductive performance in different tank designs. **(A)** Images of the housing, flat, and black-bottom flat (flat-B) tanks. **(B)** Experimental design for comparing housing and flat tanks. **(C)** Numbers of spawned eggs in housing and flat tanks, normalized to the number of females per tank to account for differences in tank design. **(D)** Fertilized egg numbers in housing and flat tanks, normalized to the total number of fish per tank. **(E)** Experimental design for comparing flat and flat-B tanks. **(F)** Fertilized egg numbers in flat and flat-B tanks. **(G)** Fertilized egg numbers in housing, flat, and flat-B tanks, normalized to the total number of fish per tank. Representative results from one of three independent trials are shown, with 4 flat and 10 housing tanks (C, D) and 3 flat and 3 flat-B tanks (F) per trial. Panel G combines the data from all three tank types shown in the preceding panels. Data are presented as box-and-whisker plots, with boxes indicating the interquartile range, center lines the median, and whiskers the minimum and maximum values. Statistical significance was determined using unpaired t-test (C, D, F) or ordinary one-way ANOVA followed by Tukey’s multiple-comparisons test (G). *p < 0.05, **p < 0.01, ***p < 0.001; NS, not significant.

To evaluate the performance of the flat tank for *D. cerebrum* breeding, we adopted the previously described spawning tube dimensions of 6 mm in inner diameter (Lam, 2022) and 5 cm in length (Schulze et al., 2018). Adult male and female fish were housed at a 1:1 ratio in either the standard housing tank (8 fish; 1.2 L) or the flat tank (20 fish; 3 L), resulting in nearly identical fish densities between the two tank types (Fig. 1B). Spawning tubes were provided at a tube-to-fish ratio of 1:2, with 4 tubes in each standard housing tank and 10 tubes in each flat tank (Fig. 1B). Following the acclimatization period, egg production and fertilization rates were assessed daily for 14 days (Fig. 1B). Fish housed in the flat tank produced significantly more eggs than those housed in the housing tank (Fig. 1C, D), indicating that the flat tank improves the reproductive performance of *D. cerebrum*.

Scototaxis, an innate preference for dark environments, has been reported in zebrafish and other teleost species (Serra et al., 1999; Maximino et al., 2010a; Maximino et al., 2010b; Blaser and Rosemberg, 2012). To determine whether a dark background influences the reproductive performance of *D. cerebrum*, we modified the flat tank by replacing its transparent bottom with a matte black bottom (Fig. 1A) and assessed egg production and fertilization rates under the same experimental conditions as in the previous experiment (Fig. 1E). Breeding pairs housed in this black-bottom flat tank (hereafter referred to as the flat-B tank) produced significantly more eggs (Fig. 1F) than those housed in the otherwise identical flat tank with a transparent bottom, yielding the highest reproductive performance among the three tank configurations tested (Fig. 1G). Together, these data indicate that optimizing spawning environments by reducing water depth, increasing horizontal space, and providing a dark background enhances the reproductive performance of *D. cerebrum*.

### Optimization of spawning tube parameters

Silicone tubes serve as effective spawning substrates for *D. cerebrum*; however, the effects of tube configurations on reproductive performance have not been systematically evaluated. To determine the optimal tube diameter, we fixed the tube length at 5 cm and the tube-to-fish ratio at 1:2 (Fig. 2A) while testing tube inner diameters ranging from 6 to 3 mm in 1-mm decrements (Fig. 2B). Following the acclimatization period, spawning tubes of each diameter (10 tubes) were placed in the flat-B tanks containing 20 adult fish (10 males and 10 females), with flat-B tanks lacking spawning tubes serving as the control (Fig. 2A), and reproductive performance was assessed once daily for 14 days (Fig. 2A).

**Figure 2.**
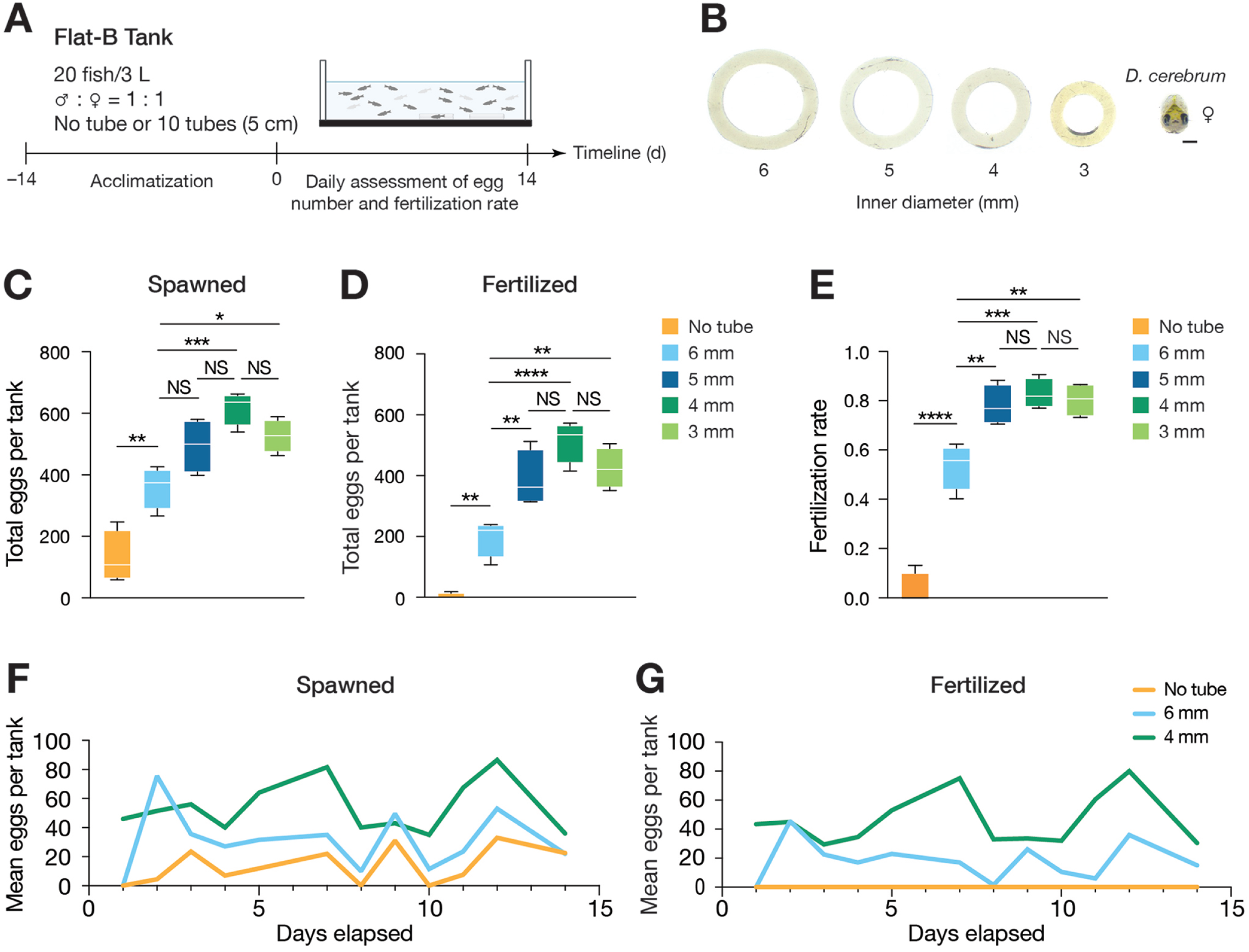
Reproductive performance with different tube diameters. **(A)** Experimental design for comparing tube diameters. **(B)** Cross-sectional images of tubes with different diameters relative to the size of an adult female. **(C–E)** Numbers of spawned (C) and fertilized (D) eggs, and fertilization rates (E), with tubes of different diameters or without tubes. **(F, G)** Trends in spawned (F) and fertilized (G) egg numbers with tubes of different diameters over the assessment period. Representative results from one of two independent trials are shown, with 2 tanks per condition (No tube, 6 mm, 5 mm, 4 mm, and 3 mm) in each trial. Data are presented as box-and-whisker plots (C–E) or as mean numbers of spawned (F) and fertilized (G) eggs over time. Statistical significance was determined using one-way ANOVA followed by Tukey’s multiple-comparisons test (C–E). *p < 0.05, **p < 0.01, ***p < 0.001, ****p < 0.0001; NS, not significant. Scale bar, 1 mm (B).

Among the tube sizes tested, 4-mm-diameter spawning tubes yielded the highest egg production (Fig. 2C, D) and fertilization rates (Fig. 2E) and consistently outperformed all other tube sizes throughout the assessment period (Fig. 2F, G). Breeding pairs, particularly those involving larger females, occasionally became lodged in 3-mm-diameter tubes, suggesting that excessive confinement may interfere with successful mating. The increase in fertilization rates with decreasing tube diameter (Fig. 2E), coupled with the observation that virtually no eggs were fertilized in the absence of spawning tubes (Fig. 2D) despite limited egg production (Fig. 2C), indicates that successful fertilization in *D. cerebrum* requires sufficient spatial confinement to ensure close proximity between breeding pairs, thereby promoting immediate contact between released sperm and eggs.

Next, we investigated the effect of spawning tube length with the tube diameter fixed at 4 mm (Fig. 3A). Tube length was varied from the standard 5 cm to 2.5 cm or 10 cm (Fig. 3B), and reproductive performance was assessed under the same experimental conditions as in the previous experiments (Fig. 3A). Egg production tended to increase with tube length (Fig. 3C, D), whereas fertilization rates remained unchanged (Fig. 3E), further supporting the importance of tube diameter in determining fertilization efficiency. Although only the difference between the 2.5- and 10-cm tubes reached statistical significance (Fig. 3C, D), the 10-cm spawning tubes consistently tended to yield higher egg production throughout the assessment period (Fig. 3F, G), suggesting that increased tube length may contribute to improved reproductive output under these breeding conditions.

**Figure 3.**
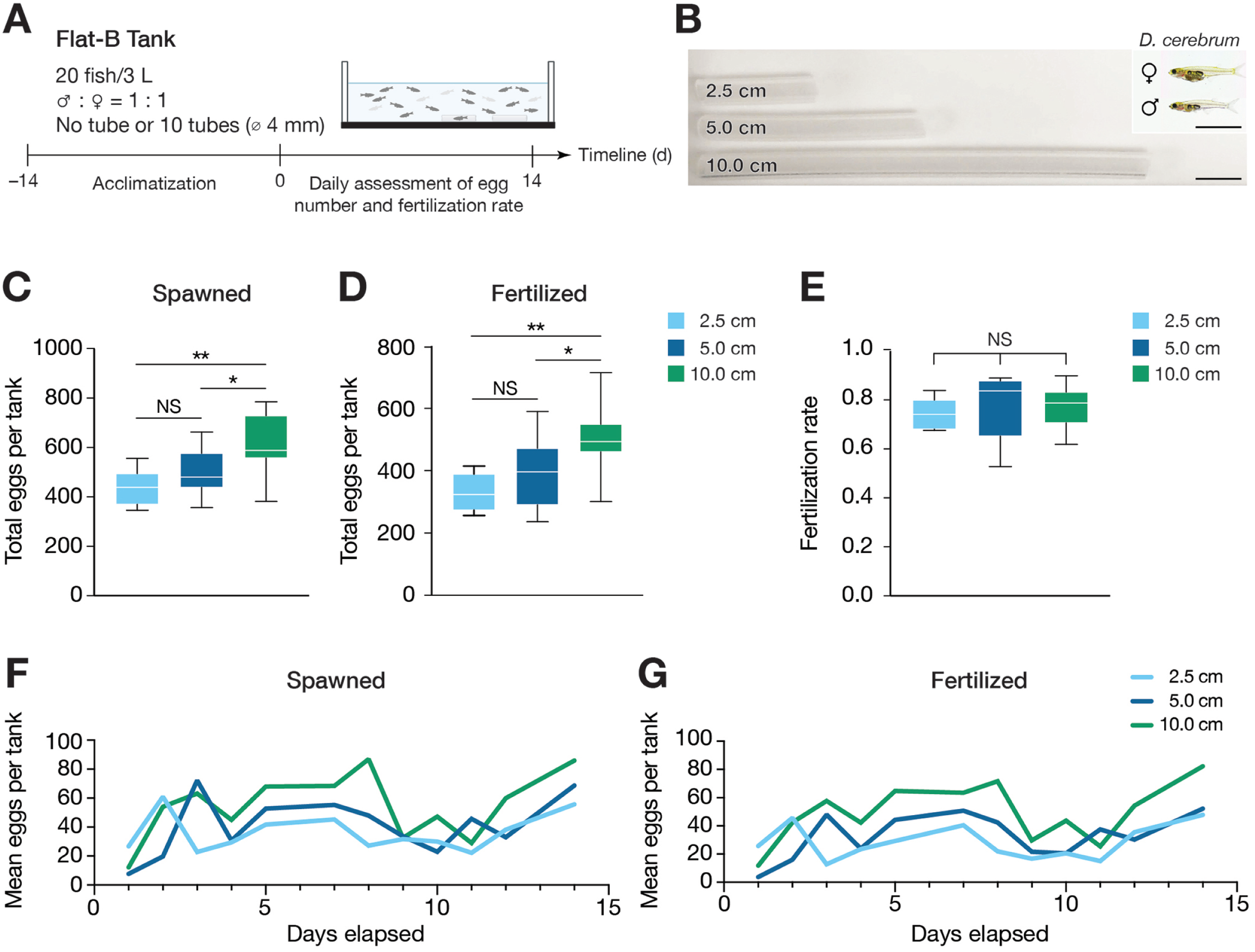
Reproductive performance with different tube lengths. **(A)** Experimental design for comparing tube lengths. **(B)** Lateral view images of tubes with different lengths relative to the size of an adult female and male. (**C–E**) Numbers of spawned (C) and fertilized (D) eggs, and fertilization rates (E), with tubes of different lengths or without tubes. **(F, G)** Trends in spawned (F) and fertilized (G) egg numbers with tubes of different lengths over the assessment period. Representative results from one of three independent trials are shown, with 3–4 tanks per condition (2.5, 5.0, and 10.0 cm) in each trial. Data are presented as box-and-whisker plots (C–E) or as mean numbers of spawned (F) and fertilized (G) eggs over time. Statistical significance was determined using one-way ANOVA followed by Tukey’s multiple-comparisons test (C–E). *p < 0.05, **p < 0.01; NS, not significant. Scale bar, 1 cm (B).

### Short-term in-tube embryo rearing

Despite a trend toward higher egg production (Fig. 3F, G), 10-cm tubes frequently retained clutches inside, making egg collection less efficient, whereas 5-cm tubes allowed clutches to detach spontaneously or be readily recovered by gentle flushing with a plastic pipette. These observations, together with the lack of a significant difference in reproductive output between the 5- and 10-cm tubes (Fig. 3C–E), support the use of 5-cm tubes as the more practical option for routine breeding. However, the tendency of 10-cm tubes to yield higher egg production while retaining egg clutches inside could be leveraged to improve workflow efficiency (Fig. 4A) when only a limited number of breeding pairs are used, such as during germline screening of individual potential founder pairs.

**Figure 4.**
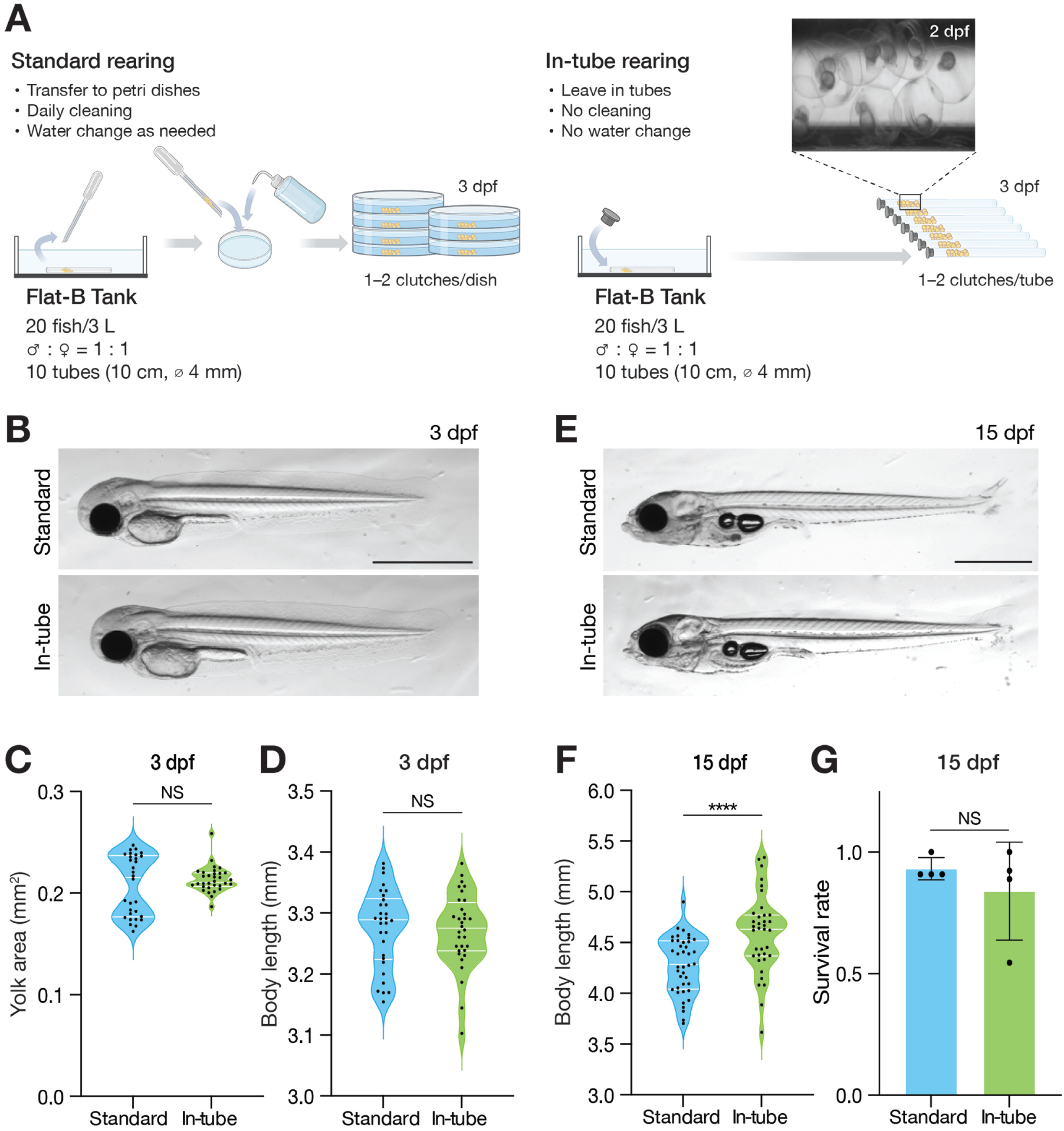
Embryonic development during short-term in-tube culture. **(A)** Workflow for standard and in-tube culture. Representative image of in-tube embryos shown in the inset. **(B)** Gross morphology of embryos following standard or in-tube culture at 3 dpf. **(C, D)** Quantification of yolk area (C) and body length (D) of embryos following standard or in-tube culture at 3 dpf (n = 30 for standard culture; n = 32 for in-tube culture). **(E)** Gross morphology of larvae at 15 dpf following 3 days of standard or in-tube culture and subsequent rearing in housing tanks. **(F, G)** Quantification of body length (F) and survival rate (G) at 15 dpf following 3 days of standard or in-tube culture and subsequent rearing in housing tanks (n = 40 for standard culture; n = 37 for in-tube culture). Data are presented as violin plots, with widths indicating data density, center lines the median, and outer lines the first and third quartiles (C, D, F), or as mean ± s.d. (G). Statistical significance was determined using unpaired t-test. ****p < 0.0001; NS, not significant. Scale bar, 1 mm (B, E). dpf, days post-fertilization.

To assess the feasibility of this approach, we collected 10-cm tubes containing one or two freshly spawned egg clutches and allowed the embryos to develop within the tubes until 3 dpf (In-tube rearing; Fig. 4A), an initial time point suitable for fluorescence screening. As a control, we collected eggs from the tubes immediately after spawning and maintained them in plastic dishes until 3 dpf using the standard rearing and maintenance protocol, including daily cleaning and/or water changes (Standard rearing; Fig. 4A). Between the in-tube and control rearing conditions, embryonic development appeared comparable through 3 dpf (Fig. 4B), and this observation was further substantiated by quantitative analysis showing no significant differences in yolk area (Fig. 4C) or body length (standard length) (Fig. 4D), established morphometric indicators of embryonic development in zebrafish (Schwartz et al., 2021).

To determine whether in-tube rearing could cause delayed adverse effects, embryos reared under both conditions until 3 dpf were subsequently maintained in housing tanks containing approximately 500 mL of system water under the same standard rearing and maintenance protocol, followed by quantification of body length and survival rate at 15 dpf. Larval development appeared comparable between groups through 15 dpf (Fig. 4E). Larval fish initially reared in tubes exhibited a modest increase in body length (Fig. 4F), with no significant difference in survival over the assessment period (Fig. 4G), indicating that short-term in-tube rearing did not adversely affect subsequent development or survival. Together, these findings suggest that in-tube rearing provides a practical approach for the short-term maintenance of individual clutches, offering improved space efficiency and reduced handling for applications such as transgenic line propagation.

### Spawning tube selection during mating

In addition to tube diameter and length, we varied the number of spawning tubes with the expectation that greater availability of spawning sites would promote egg production (Fig. S1A). However, reproductive performance was unaffected by a 4-fold difference in spawning site availability (Fig. S1B), raising a possibility that breeding pairs may preferentially reuse the same tubes despite the availability of alternative spawning sites.

To address this possibility, we individually labeled each spawning tube (Fig. 5A) and monitored tube usage over 10 days by recording the numbers of clutches and eggs deposited in each tube daily (Fig. 5B). We used 15-cm-long tubes to maximize clutch retention (arrows; Fig. 5B), thereby enabling accurate counting of clutches and eggs within individual tubes. The presence of multiple clutches within a single tube was considered indicative of repeated use by different breeding pairs, as *D. cerebrum* females typically spawn no more than once per day (Rajan et al., 2022a). Although some eggs became dislodged from the spawning tubes, the majority remained within the tubes during the assessment period (Fig. 5C), supporting reliable tracking of clutch deposition. Moreover, an average of 2–4 clutches were produced per tank each day (Fig. 5D), confirming that multiple breeding events occurred daily under these experimental conditions.

**Figure 5.**
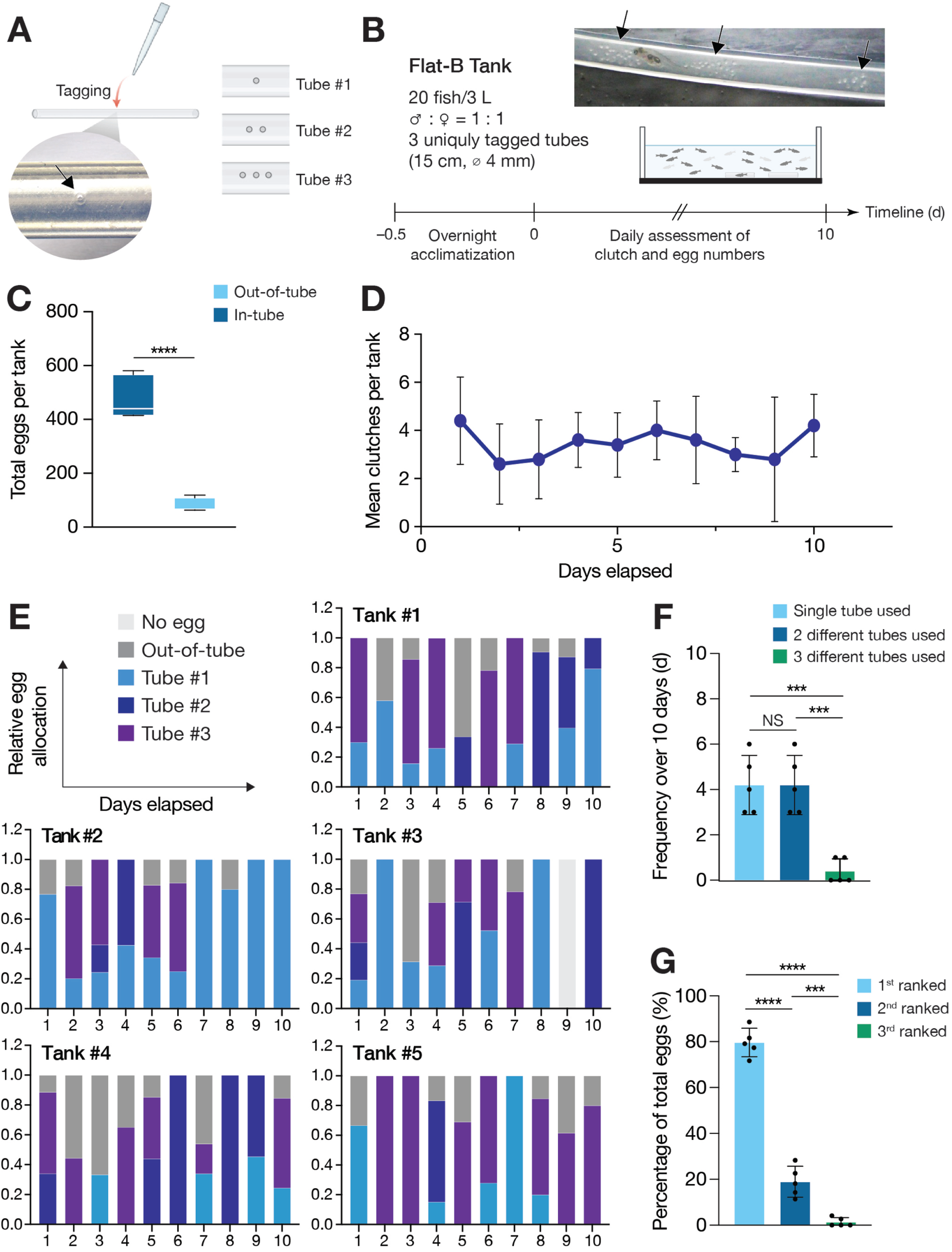
Analysis of spawning tube selection. **(A)** Tagging of individual spawning tubes with unique embossed patterns (arrow). **(B)** Experimental design for assessing spawning-site selection. Representative image of multiple clutches (arrows) within a tube shown in the inset, indicating multiple mating events. **(C)** Numbers of eggs spawned inside and outside the tubes per tank over the assessment period. **(D)** Clutch numbers over the assessment period. Data are shown as the mean of five independent tanks. **(E)** Daily patterns of tube selection over the assessment period across all five tanks. **(F)** Frequency of days on which clutches were deposited in one, two, or three different tubes during the 10-day assessment period. **(G)** Percentage of total eggs deposited in each tube according to its frequency of use. Data are presented as box-and-whisker plots (C), mean ± s.d. (D, F, G), or column charts (E). Five tanks were analyzed (C–G). Statistical significance was determined using Unpaired t-test (C) and ordinary one-way ANOVA followed by Tukey’s multiple-comparisons test (F). ***p < 0.001, ****p < 0.0001; NS, not significant.

We next determined which spawning tubes were used for breeding events in each tank. Strikingly, spawning was confined to single or two different tubes per day in nearly all cases, with three tubes used on only two occasions (Day 3 in Tank 2 and Day 1 in Tank 3; Fig. 5E). The most frequently used tube varied from day to day within each tank (Fig. 5E), indicating no consistent preference for a particular spawning tube. Quantification of daily tube usage showed that spawning in a single tube and in two different tubes occurred at comparable frequencies (Fig. 5F). Despite this variation in tube usage, quantification of egg distribution among tubes revealed a pronounced bias toward a single tube (Fig. 5G), providing evidence that *D. cerebrum* breeding pairs preferentially reuse spawning sites previously used by other pairs.

## Discussion

In this study, we systematically evaluated tank configurations and spawning-tube parameters to improve egg production and collection efficiency in *D. cerebrum*. Our results show that the use of a black-bottom tank under otherwise identical conditions enhances egg production, identifying the visual characteristics of the spawning environment as an important factor influencing reproductive output. Among spawning-tube parameters, tube diameter had a major influence on reproductive performance, with narrower tubes increasing both egg production and fertilization efficiency and a 4-mm inner diameter yielding the highest egg production. Tube length also influenced reproductive performance, with longer tubes showing a trend toward increased egg production. For routine egg collection, 5-cm tubes are more practical because clutches can be recovered more readily. In contrast, the tendency toward higher egg production and greater clutch retention in 10-cm tubes may be advantageous when frequent egg collection is unnecessary, as supported by our demonstration that embryos can be maintained directly within the tubes for short-term culture. Lastly, we showed that *D. cerebrum* exhibits a preference for previously used spawning sites, rather than simply responding to increased spawning site availability.

Spawning-tube diameter strongly influenced both egg production and fertilization efficiency in *D. cerebrum* (Fig. 2C–E). The greater reproductive output observed with decreasing tube diameter is likely attributable to the confined space provided by narrower tubes, which may facilitate the close physical interaction between mating pairs required for successful spawning. Consistent with this possibility, spawning events were markedly reduced in 6-mm tubes (Fig. 2C), suggesting that tube dimensions themselves influence spawning behavior. Spatial confinement also appears critical for fertilization, as it would restrict sperm dispersal and maintain a higher local sperm concentration around freshly released eggs. Indeed, the few eggs spawned in 6-mm tubes remained virtually unfertilized (Fig. 2C, D), supporting the importance of sufficient confinement for successful fertilization. Together, these findings demonstrate that *D. cerebrum* mating behavior is favored by a confined physical environment, experimentally recapitulated by the 4-mm tubes used in this study. This preference may reflect adaptation to similarly confined structures encountered in natural spawning habitats, although the physical characteristics of such sites remain unknown.

Notably, increasing the number of available spawning tubes did not further increase egg production (Fig. S1B), and the analysis of tube usage revealed a preference for spawning sites that had already been used (Fig. 5G). A similar spawning-site preference has been reported in bluefin killifish, *Lucania goodei*, a non-nest-building species that lacks parental care. Female *L. goodei* preferentially deposit eggs at sites already containing conspecific eggs, a behavior proposed to enhance offspring survival either through a dilution effect, whereby clustering eggs with those of other females reduces the predation risk for each individual egg, or by selecting sites favorable for offspring development (Welsh and Fuller, 2011). *D. cerebrum*, another non-nest-building species lacking parental care, may employ a similar strategy to enhance offspring survival. Initial spawning-site selection appeared relatively variable (Fig. 5F), whereas subsequent spawning became concentrated in a single tube (Fig. 5G), suggesting that early spawning events may guide subsequent selection of a preferred spawning site. The cues underlying this selection remain unknown but could involve visual or chemical signals associated with prior spawning activity or deposited eggs. In this context, the tendency toward greater egg production in 10-cm tubes may be related to their greater retention of deposited eggs, which could provide more persistent visual and/or chemical cues for subsequent spawning. Conversely, 2.5-cm tubes yielded fewer eggs despite appearing to provide sufficient space for mating (Fig. 3B), possibly because eggs were more readily dislodged and therefore less able to provide such cues. Given that *D. cerebrum* is a highly social species in which males exhibit vocal communication (Schulze et al., 2018), it is intriguing to consider whether acoustic signals contribute to communication among individuals during spawning-site selection.

## Acknowledgements

We thank Dr. Benjamin Judkewitz, Charité–Universitätsmedizin Berlin, Germany, for providing *D. cerebrum* and advice on its husbandry. We also thank Dr. Naoki Mochizuki, BIKEN Foundation, Japan, for support and advice, and Yasuko Morita, Shigemi Toyoshima, Tomoe Sato, Yoshimi Ando, Maria Fujimoto, Rei Yanagida, Naoko Kobayashi, and Minami Fujita for fish husbandry and animal care at the NCVC fish facility. Figures were prepared in part using BioRender.com (Kao, C. C., 2027; https://BioRender.com/n9mdznk).

## Author contributions

Conceptualization: C.C.K., K.A., K.K.; Analysis: C.C.K., D.S.J.; Funding acquisition: K.K.; Project administration: T.K., K.K; Writing–original draft: C.C.K., K.K.; Writing– review & editing: C.C.K., D.S.J., K.A., T.K., K.K.

---

**Figure S1.**
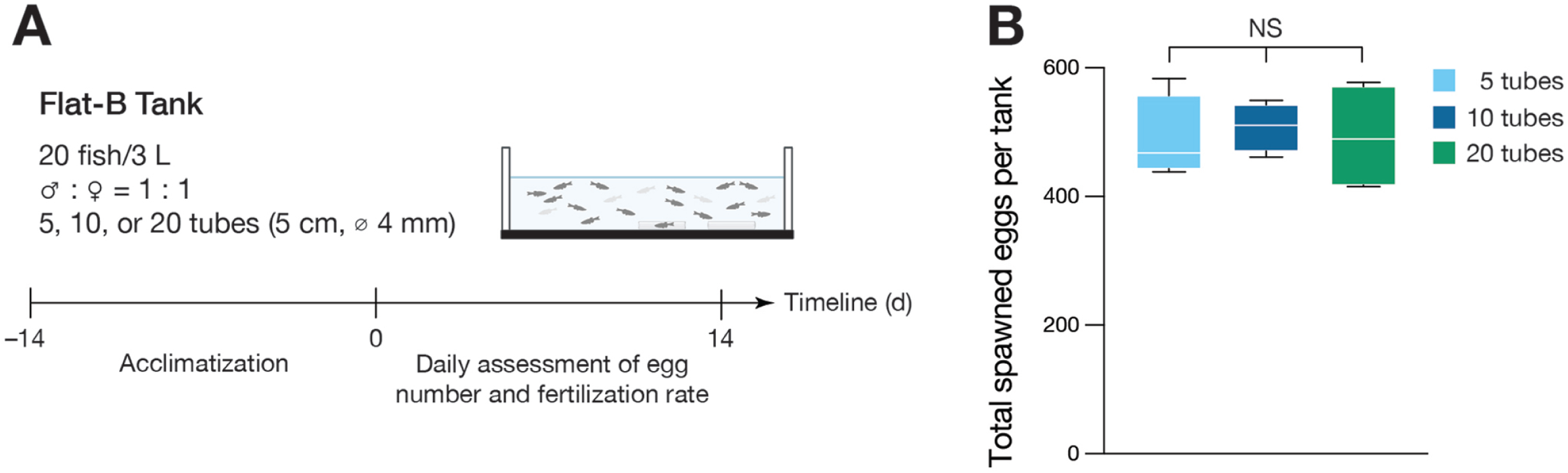
Reproductive performance with different numbers of spawning tubes. **(A)** Experimental design for comparing egg production with different numbers of spawning tubes. **(B)** Numbers of total spawned eggs per tank with different number of tubes. Fertilization rates were comparable across conditions. Statistical significance was determined using one-way ANOVA followed by Tukey’s multiple-comparisons test (B). Data are presented as box-and-whisker plots, with four tanks analyzed per condition. NS, not significant.

